# The “abominable mystery” of Schenck: the polymorphism of *Serjania piscatoria* and its implications for the evolution of vascular variants in Paullinieae (Sapindaceae)

**DOI:** 10.1101/2024.04.04.587907

**Authors:** Natália F. Marques, Israel L. Cunha Neto, Lilian A. Brito, Genise V. Somner

## Abstract

*Serjania* is the only genus of Paullinieae that exhibits all types of vascular variants in stems and includes *S. piscatoria* with a complex vascular structure that has intrigued botanists for centuries. Here, we analyzed the stem development of *S. piscatoria* in an evolutionary context and determined its phylogenetic position within the genus. We studied four individuals using standardized anatomical techniques and employed DNA sequencing and phylogenetic analysis to determine the species’ phylogenetic position. Additionally, we employed ancestral state reconstruction to explore the pattern of evolution of vascular variants. We find that the stem development in *S. piscatoria* is determined by various ontogenetic processes that result in vascular variants that occur through modifications during primary and/or secondary growth, or ectopic cambia formation. These various patterns are classified into distinct categories of vascular variants, highlighting the lability of vascular meristems and the polymorphism within the species, which manifests across different individuals. *Serjania piscatoria* belongs to a clade composed of species with compound stems, from which the fissured stems observed in the species would have evolved. The findings provide evidence for the diverse stem vasculature in *Serjania*, and the importance of studying vascular variant diversity from a developmental and evolutionary perspective.

---

> “*Serjania piscatoria* Radlk. stands out among all species of the genus, as far as I know, by the most complex anomalies in its older stems.” Heinrich Schenck (1893, p.100; translation from original in German)

## INTRODUCTION

The ancestor of seed plants is hypothetically characterized as having a stem development formed by the combination of an eustele – a single ring of vascular bundles – and the formation of secondary xylem and phloem produced by a single bifacial cambium (Spicer and Groover, 2010; Ragni and Greb, 2018; Onyenedum and Pace, 2021). While this typical development results in stems with homogeneous amounts of wood (secondary xylem) and inner bark (secondary phloem) in most extant seed plants, many lineages have evolved alternative developmental pathways (=ontogeny) that deviate from this typical vascular architecture. These alternative ontogenies lead to anatomical modifications in the rate and/or organization of vascular tissues due to changes in the location, organization, and activity of vascular meristems in the plant axis and are here referred to as *vascular variants* (previously known as “anomalous growth” or “cambial variants”; Cunha Neto, 2023). Besides generating considerable morphological diversity, the study of vascular variants has gained notoriety due to their adaptive roles in climbing plants (e.g., flexibility, conductivity, injury repair, and mechanical resistance; Carlquist, 1991, 2013; Fisher and Ewers, 1991; Rowe et al., 2004) and self-supporting plants including evidence from mangrove trees (e.g., adaptation to dynamic mangrove environment; Robert et al., 2014) or herbaceous species (e.g., alternative routes for increased hydraulic conductivity through medullary bundles; Cunha Neto et al., 2023). Additionally, different types of vascular variants generating such functional forms may have evolved in a correlated fashion and can be associated with species diversification (Cunha Neto et al., 2022, 2023). Ultimately, the unique patterns resulting from the establishment of vascular variants provide important information for the systematics of groups containing lianas (i.e., woody vines) (Carlquist, 2001; Pace et al, 2009; Angyalossy et al., 2012, 2015; Acevedo-Rodriguez et al., 2015 (onwards)), and their stems with such intriguing forms are a source of material for different artistic projects in tropical countries (e.g., marchetery; Tamaio and Somner 2010). From an evolutionary standpoint, understanding the evolution of development of vascular variants is crucial to reveal the exact ontogenetic processes that determine how these forms originated and diversified through evolutionary time. Consequently, elucidating plant diversity and evolution requires knowledge of the anatomy and development of plants with such complex anatomies, which has received well-deserved attention from botanists worldwide.

Since the seminal work of Ludwig Radlkofer (1875) and Heinrich Schenck (1893), significant emphasis has been placed on investigating the anatomy, development, and evolution of vascular variants in Sapindaceae, as this family has been recognized as having the greatest diversity of anatomical patterns across seed plants (Araújo and Costa 2006; Tamaio and Angyalossy 2009; Cunha Neto et al., 2018; Chery et al., 2020; Rajput et al., 2021; Rizzieri et al., 2021; Pace et al., 2022). Currently, in addition to the typical growth observed in stems of seed plants (i.e., eustele + single bifacial cambium), ten types of vascular variants are recognized for the family (Pace et al., 2022). In this classification, of the ten vascular variants recognized for Sapindaceae, nine are recorded for *Serjania*, except for the “fissured vascular cylinder” (fissured stems) (Pace et al., 2022). Originally, Schenck (1893) grouped the species *Serjania piscatoria* Radlk. with the genus *Urvillea* in the same “fissured stem” category. According to the author, the fissured vascular cylinder has two distinct origins: in *S. piscatoria*, it originates with the formation of phloem wedges interrupting the secondary xylem, while the stem of *Urvillea* begins with a lobed stem outline in primary growth, and subsequent fissuring of stem portions occurs through the tissue expansion that reaches the pith and can “fissure” the stem into two or more independent units. As indicated by Schenck (1893) and others (Pace et al., 2022), different developmental processes underlie the “fissured stems” in *Urvillea* and *S. piscatoria*. Recently, Cunha Neto et al. (2023) reported five developmental pathways in *Urvillea* stems, with only one ontogeny (exclusive to the species *U. laevis* Radlk.) exhibiting a developmental pathway similar to the “fissured” stem of *S. piscatoria*. While *U. laevis* and *S. piscatoria* share a similar pattern of fragmentation of the vascular system (without stem splitting), stem split into independent units is observed only in other *Urvillea* species and results from different ontogenetic processes (Cunha Neto et al. 2023). In contrast, ectopic cambia are frequently observed in *S. piscatoria* (Schenck 1893; Bastos et al., 2016) but were reported only for *U. filipes* Radlk. (Cunha Neto et al. 2023), and additional anatomical modifications seem to be present in *S. piscatoria* (Fig. 1A-E; Schenck, 1893). Studies on the evolution of vascular variants in *Urvillea* support the notion that there are significant differences between the vascular organization observed in *S. piscatoria*, indicating the need for a deeper examination of its developmental diversity.

**Figure 1.**
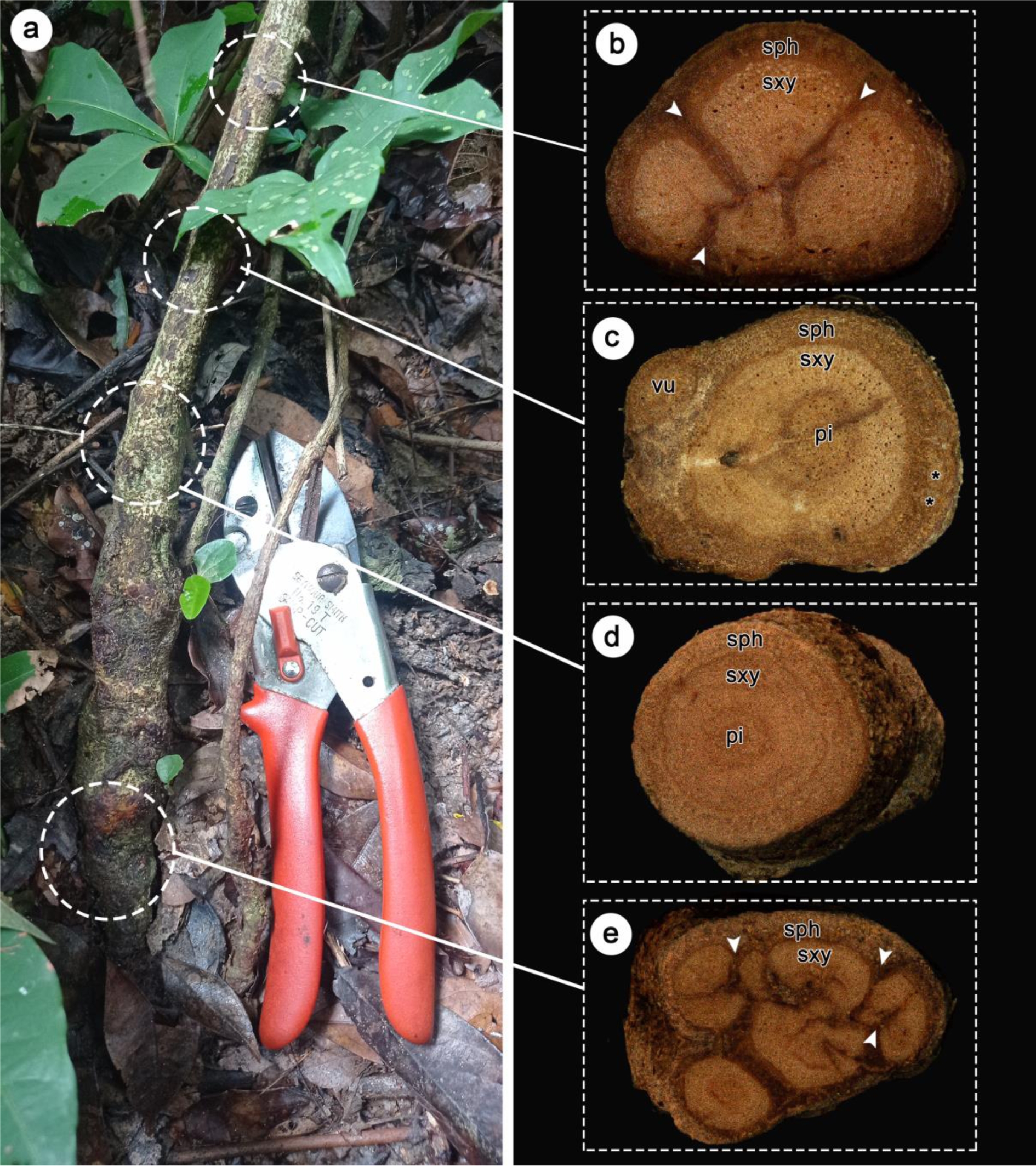
External morphology and macroscopic view of stems of *Serjania piscatoria* (individual 2). A, Basal portion of the plant. B–E, Stem cross-sections sampled 150–200 mm away from the base of the plant of A, showing vascular system compartmentalized by deep phloem wedges (B, white arrowheads), isolated vascular unit (vu) derived from an isolated vascular bundle and neoformations (ectopic cambia) (C, asterisks). D, Stem with typical secondary growth. E, Phloem wedges fragmenting the vascular system (white arrowheads). Images were photographed at intervals of approximately 50 mm. Abbreviations: sxy = secondary xylem, pi = pith, sph = secondary phloem. Stem diameter: B = 14 mm, C = 20 mm, D = 16 mm, E = 22 mm.

The phylogeny of *Serjania* by Steinmann et al. (2022) is the first to attempt to expand the phylogenetic sampling of this lineage that was previously included only in tribe-level and family-level phylogenies. Out of nearly 250 species encompassing the genus, this study included only 43 species, focusing on species with unusual fruits and delimitation of new species from Mexico. *Serjania piscatoria*, the focus of this study, was not included in that phylogenetic investigation. This species is endemic to Brazil, restricted to the Atlantic Forest, and has only a few collected specimens (e.g., 107 records), even though it had been described in the late 19^th^ century by Radlkofer (1875). Surprisingly, the anatomy of *S. piscatoria* was documented in the work of Schenck (1893) with notorious notes on the complex vascular organization of the species. Yet, further developmental studies and a phylogenetic investigation including the species are still lacking. Here, we investigated the stem development of *S. piscatoria* to understand the ontogenetic processes that generate its structural diversity in a systematic and evolutionary context. In addition, we tested the phylogenetic position of *S. piscatoria* within the recent phylogeny of the genus (Steinmann et al., 2022) and the pattern of evolution of vascular variants within this same phylogenetic framework.

## MATERIALS AND METHODS

### Plant collection

Fresh specimens from four individuals of *Serjania piscatoria* were obtained from plants growing in natural populations of the Atlantic Forest in Rio de Janeiro State, Brazil (Table 1). Stem samples were systematically collected from various parts of the plants (sections of approximately 50 mm in length), from young stems (including the shoot apex) to mature stems (towards the base of the plant). Samples with smaller diameters (1-7 mm) and larger diameters (10-50 mm) were fixed in FAA (formaldehyde, glacial acetic acid, and 70% ethanol) and subsequently stored in 70% ethanol (adapted from Johansen, 1940). Voucher specimens were deposited in the herbarium of the National Museum (R) and the Federal Rural University of Rio de Janeiro (RBR). Wood samples were deposited in the wood collection of the Rio de Janeiro Botanical Garden (RBw) and the National Museum (Rw).

**Table 1.**
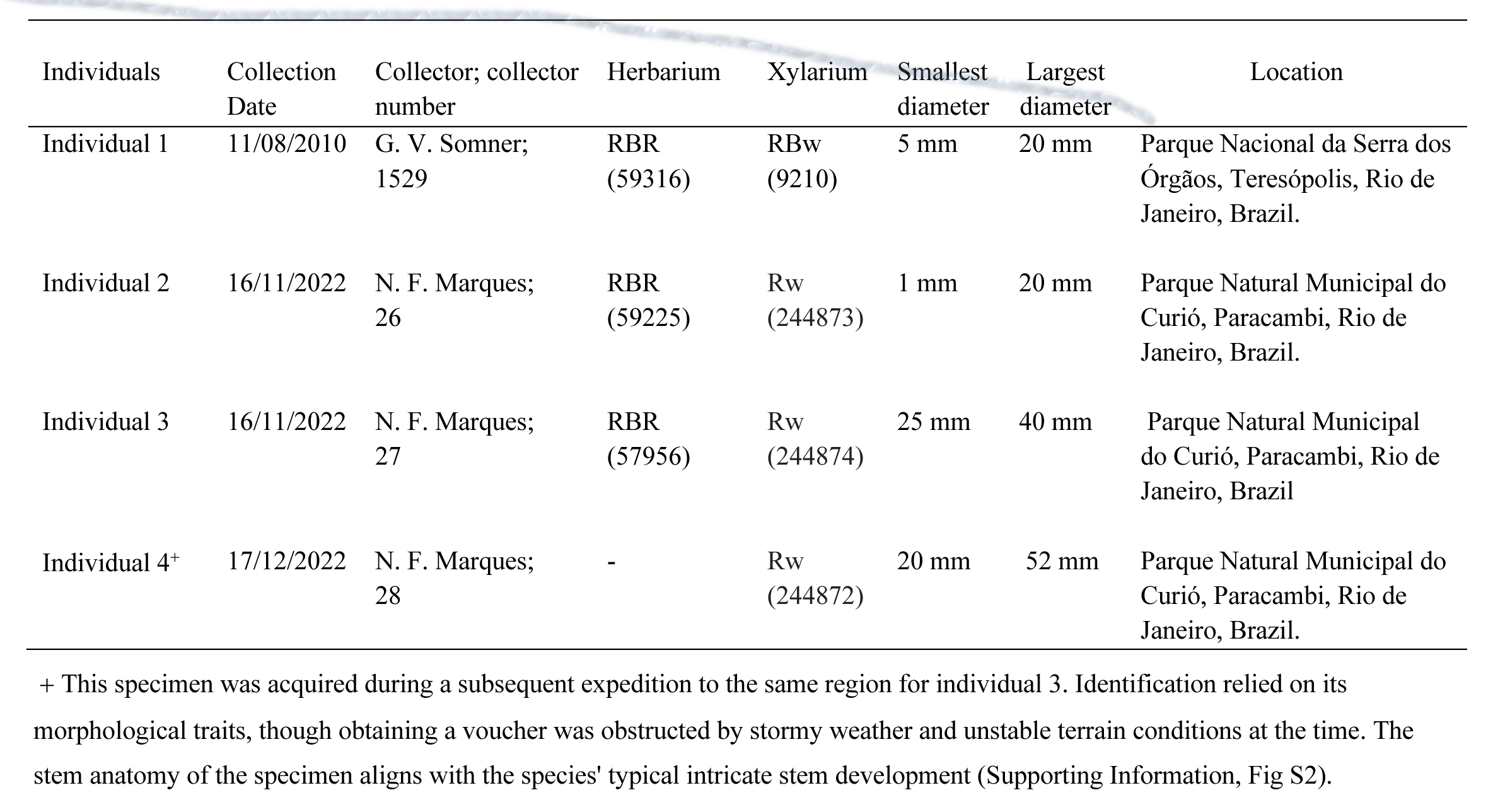
List of studied specimens of *Serjania piscatoria* Radlk., including information on collection data, accession number, and stem diameter. All specimens were collected in natural areas of the Atlantic Forest in Rio de Janeiro, Brazil.

| Individuals | Collection Date | Collector; collector number | Herbarium | Xylarium | Smallest diameter | Largest diameter | Location |
| --- | --- | --- | --- | --- | --- | --- | --- |
| Individual 1 | 11/08/2010 | G. V. Somner; 1529 | RBR (59316) | RBw (9210) | 5 mm | 20 mm | Parque Nacional da Serra dos Órgãos, Teresópolis, Rio de Janeiro, Brazil. |
| Individual 2 | 16/11/2022 | N. F. Marques; 26 | RBR (59225) | Rw (244873) | 1 mm | 20 mm | Parque Natural Municipal do Curió, Paracambi, Rio de Janeiro, Brazil. |
| Individual 3 | 16/11/2022 | N. F. Marques; 27 | RBR (57956) | Rw (244874) | 25 mm | 40 mm | Parque Natural Municipal do Curió, Paracambi, Rio de Janeiro, Brazil |
| Individual 4<sup>+</sup> | 17/12/2022 | N. F. Marques; 28 | - | Rw (244872) | 20 mm | 52 mm | Parque Natural Municipal do Curió, Paracambi, Rio de Janeiro, Brazil. |

### Macroscopic analyses

For macroscopic preparation (Fig. 1A-E), samples with larger diameters were polished using a series of increasingly fine waterproof sandpapers (e.g., 600, 1200, 2000), sliding the samples on the sandpaper under water (adapted from Barbosa et al., 2021). The polished samples were stored in 70% ethanol, photographed with a digital camera (Canon Eos Rebel T3I), and under a 3D microscope (ZEISS Smartzoom 5).

### Anatomical procedures for light microscopy

To detect the main developmental stages, stem samples were initially hand-sectioned using a razor blade. Some sections generated through this technique provided images for the study. Subsequently, samples with smaller diameter were dehydrated in an ethylic series (70%–100%), embedded in historesin (Historesin® – Leica), sectioned using a rotary microtome (Leica RM 2245) with glass knives (Leica), with thickness ranging from 3 to 5 µm. Staining was performed with 0.05% toluidine blue (O’Brien and McCully, 1981). Larger stem portions were softened in boiling water with glycerin and then immersed in more concentrated solutions of polyethylene glycol 1500 (10% to 100%) (Rupp, 1964). These samples were sectioned at variable thickness (approx. 16-25 µm) using a sliding microtome (Leica SM2000R), stained with 1% Astra Blue and 1% Safranin, and mounted in Entellan for the preparation of permanent slides (adapted from Barbosa et al., 2010). Photographs were taken using a Leica DM750 Microscope coupled with a Leica ICC50 HD camera and LAS EZ software version 3.0.0.

### Molecular data collection

To identify the phylogenetic relationship of *Serjania piscatoria* in *Serjania* phylogeny, the data obtained in the phylogenetic study by Steinmann et al., (2022) was used. The new phylogeny included 60 species from the tribe Paullinieae, using data from two DNA regions: ITS (nuclear) and trnL intron (chloroplast). The inner group consisted of 49 *Serjania* species, including *S. piscatoria*, and the outer group comprised 11 species from other genera of the tribe. The new alignment is available on Zenodo (https://zenodo.org/records/10821731).

### DNA extraction, amplification, and sequencing

DNA sequences were generated for *S. piscatoria* using the same regions as Steinmann et al., (2022) (i.e., ITS and trnL). Total genomic DNA was extracted following the manufacturer’s protocol “DNeasy Plant Pro Kit” from QIAGEN. For amplification, the primers ITS4 and ITS5 were used for the ITS region (White et al., 1990), and the trnL intron was amplified with primers trnl-c and trnl-d from Taberlet et al. (1991). The PCR products were subjected to Sanger sequencing using the “ABI 3730xl System” sequencer from Macrogen. The sequences were deposited in the Genbank data repository (accession numbers: PP565355 and PP573764).

### Alignment and phylogenetic analysis

*Serjania piscatoria* sequences were processed using Geneious v.8.0.5 (Bio-matters Ltd., Auckland, New Zealand and aligned with selected sequences from Steinmann et al. (2022). Geneious v.8.0.5 software was used for individual alignments of each region and the concatenated alignment of both regions, using MUSCLE with default parameters. The concatenated alignment was used for maximum likelihood (ML) and Bayesian inference (BI) phylogenetic analyses. The most suitable partitioning scheme was obtained using the IQ-TREE feature on the CIPRES Science Gateway portal (Miller et al., 2010). IQ-TREE v.2.1.2 software was used for ML analysis with 1000 bootstrap replicates. For Bayesian partitioned analysis, a modified set of substitution models (based on the ModelFinder Result) in MrBayes v.3.2.7. The Markov Chain Monte Carlo was run with two executions on each of the four chains, for 10 million generations with 25% burn-in of sampled trees every 1000 generations. Chain convergence was assessed using log posterior probability and effective sample size (≥200), analyzed by Tracer v.1.7.2 (Rambaut et al., 2014). Posterior tree probabilities were combined using BBEdit v.14.6.3 (Bare Bones Software; http://www.barebones.com/). Subsequently, TreeAnnotator v.1.10.4 (BEAST v.1.10 package; Suchard et al., 2018) was used to generate the maximum clade credibility (MCC) tree using combined MrBayes execution tree possibilities with median node heights.

### Character coding and ancestral character state analysis

To investigate the pattern of evolution of vascular variants across *Serjania*, we first determined the variant types in *S. piscatoria* and closely related species. Following previous studies on evolution of development of vascular variants for Paullinieae, Sapindaceae (Chery et al., 2020, Cunha Neto et al., 2023), each developmental trajectory was coded as a character state (Supporting Information, Dataset S1). Ancestral states were estimated and visualized using stochastic character mapping with the make.simmap function (Revell, 2013) under the best-fitting evolution model (ER) determined by the Akaike Information Criterion in the fitDiscrete function of the R package Geiger (Pennell et al., 2014). A thousand simulations were performed along the maximum likelihood tree, and results were summarized using the plot_simmap function written by Dr. Michael May (University of California, Berkeley, USA). All analyses were conducted in R (R Core Team, 2022) using a script adapted from Cunha Neto et al., (2023), available on Zenodo (https://zenodo.org/records/7754041).

## RESULTS

In *Serjania piscatoria*, the distribution of vascular bundles in most specimens assumes an irregular arrangement leading to the formation of a wavy contour of the vascular cambium originating from these bundles (Fig. 2A-C). The wavy contour of the cambium is maintained/intensified by the differential rate in the production of its derivatives in some regions of the stem circumference (Fig. 2C). Some specimens can also present isolated bundles in young stems, which may or may not give rise to a vascular unit with secondary growth in adult stems (Fig. 3A-E, 4A-E). In addition, *de novo* vascular tissues can be produced through the emergence of ectopic cambia at later developmental stages (Fig. 5A-F, 6A-H). Thus, throughout stem development, all individuals of *S. piscatoria* may exhibit one or more of the following four ontogenetic processes: (1) independent vascular units derived from isolated bundles; (2) phloem wedges derived from differential cambial activity; (3) compartmentalization of the vascular system through continuation of differential cambial activity; and (4) ectopic cambia. Below, we describe in detail each of these processes and how they individually or collectively contribute to the development of the vascular system in stems of different specimens of *S. piscatoria*.

**Figure 2.**
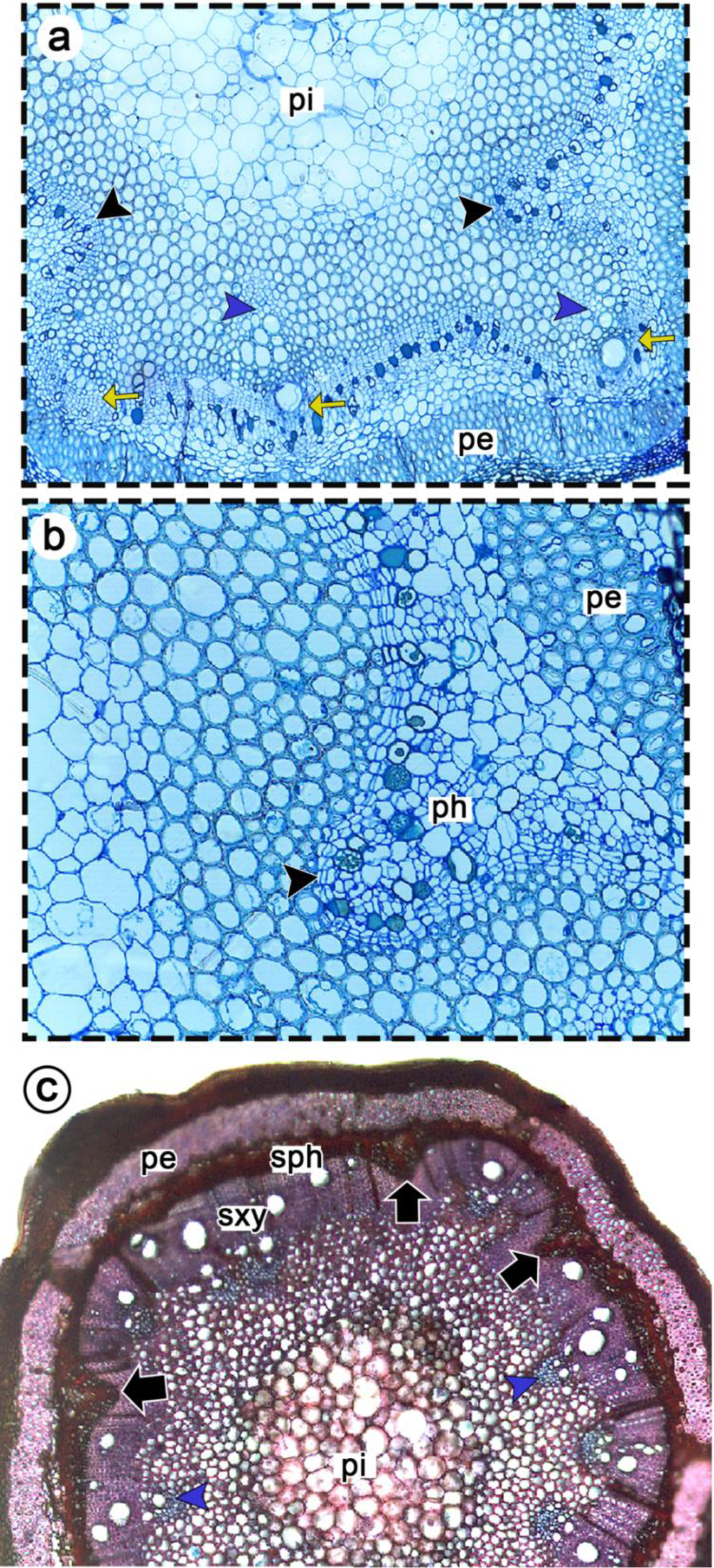
Cross sections of stems of *Serjania piscatoria* during transition from primary to secondary growth. A, B Individual 1. Developing cambium with wavy contour due to asymmetrical distribution of vascular bundles and differential activity in the production of xylem and phloem (A), mostly in the region of the interfascicular cambium (black arrowhead; detail in B). The yellow arrows indicate the fascicular cambium. C, Individual 2. Initial secondary growth with the formation of phloem wedges (black arrows) in regions of the interfascicular cambium. blue arrowheads = protoxylem poles, pe = fibrous pericycle, ph = phloem; sxy= secondary xylem; sph= secondary phloem.

**Figure 3.**
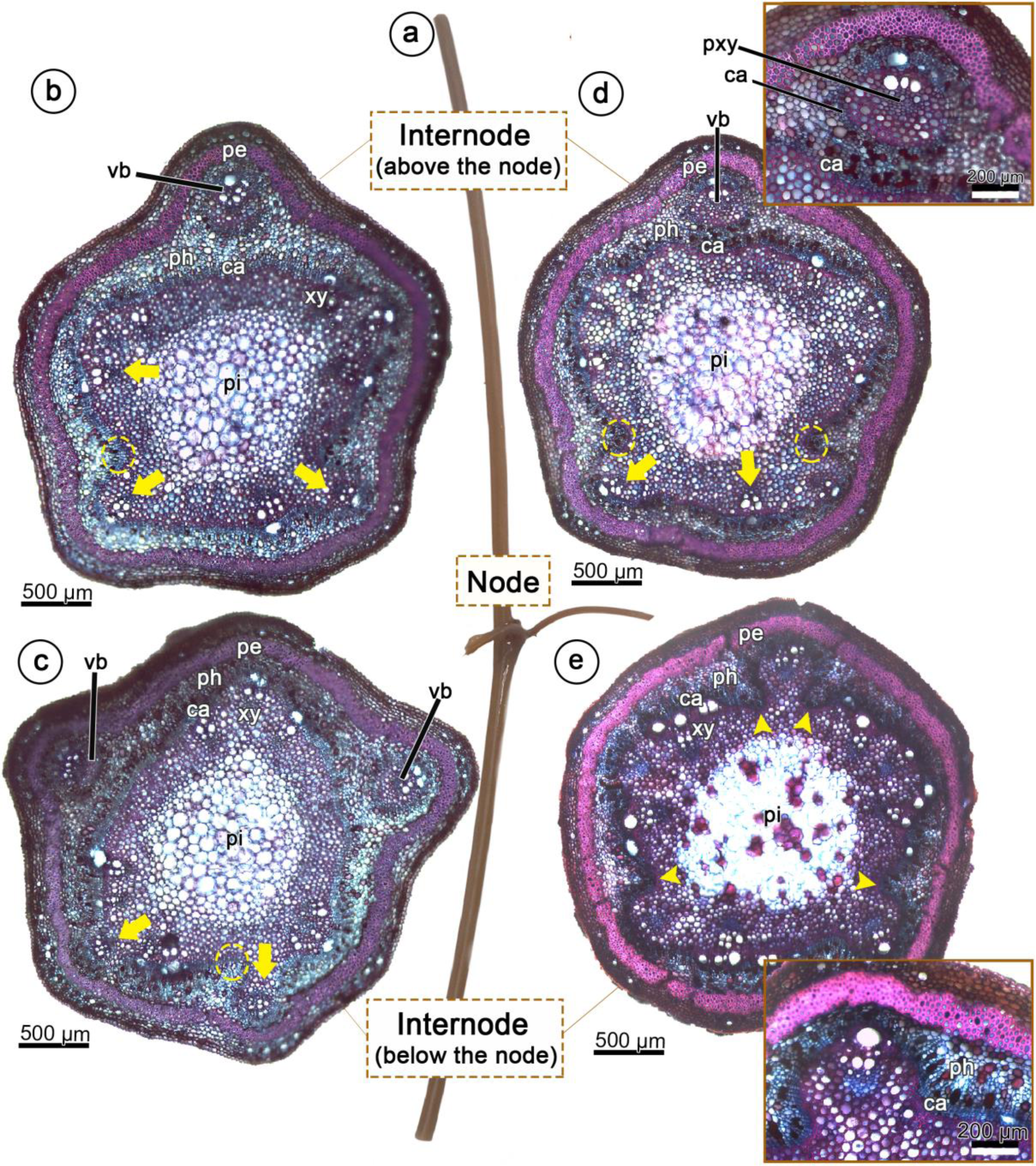
Young stem of *Serjania piscatoria* with presence or absence of an isolated vascular bundle (individual 2). A, Stem with two developed internodes. B–E, Cambium development and initial phloem arcs (dotted yellow ellipse); note the presence of one or two isolated vascular bundles (vb), and a displaced vascular bundle connected with the central cylinder through the developing cambium in E (inset), which forms phloem arcs (yellow arrowheads). The yellow arrows point to the relative position of the collateral vascular bundles. B–C Specimen 1 mm thick. D–E Specimen 2 mm thick. Abbreviations: ca = developing cambium, pe = fibrous pericycle, ph = phloem, pi = pith, vb = vascular bundle, xy = xylem.

**Figure 4.**
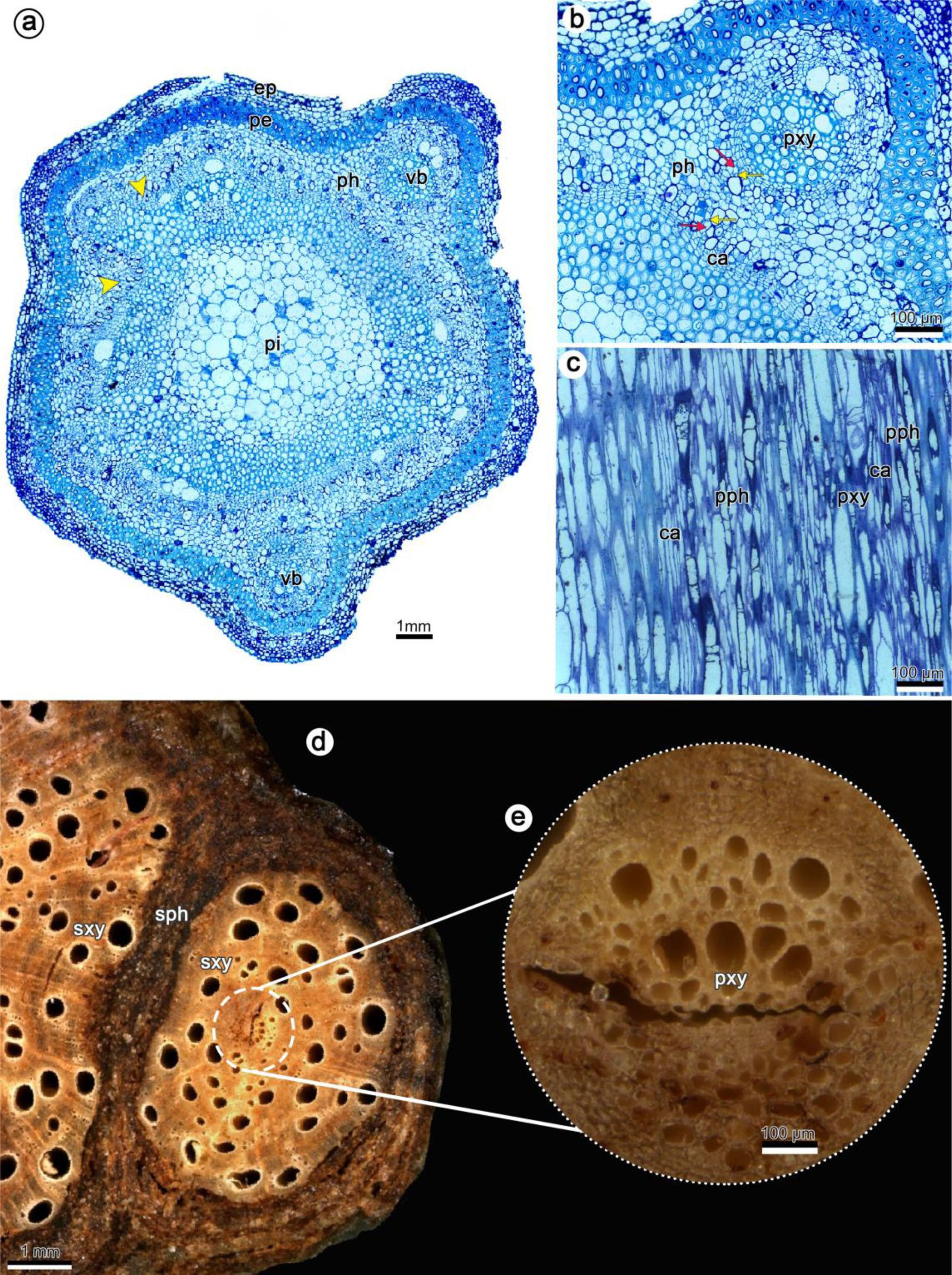
Development of compound structure in stems of *Serjania piscatoria* (individual 2). A, Note isolated vascular bundles (vb) and arcs of phloem (yellow arrowheads). B–C, Detail of A showing isolated vascular bundle in cross-section (B) and longitudinal view (C); note primary phloem with sieve tube elements (red arrow) and companion cells (yellow arrow) between the bundles of eustele and the isolated bundle in B. D–E, Macroscopic view showing secondary vascular tissues forming the “peripheral vascular cylinder” derived from the isolated vascular bundle, which is evidenced from the presence of primary xylem. Abbreviations: ep = epidermis, pe = pericycle, vb = isolated vascular bundle, pxy = primary xylem, pc= procambium, pi = pith, pph = primary phloem, pxy = primary xylem, sxy = secondary xylem, sph = secondary phloem.

**Figure 5.**
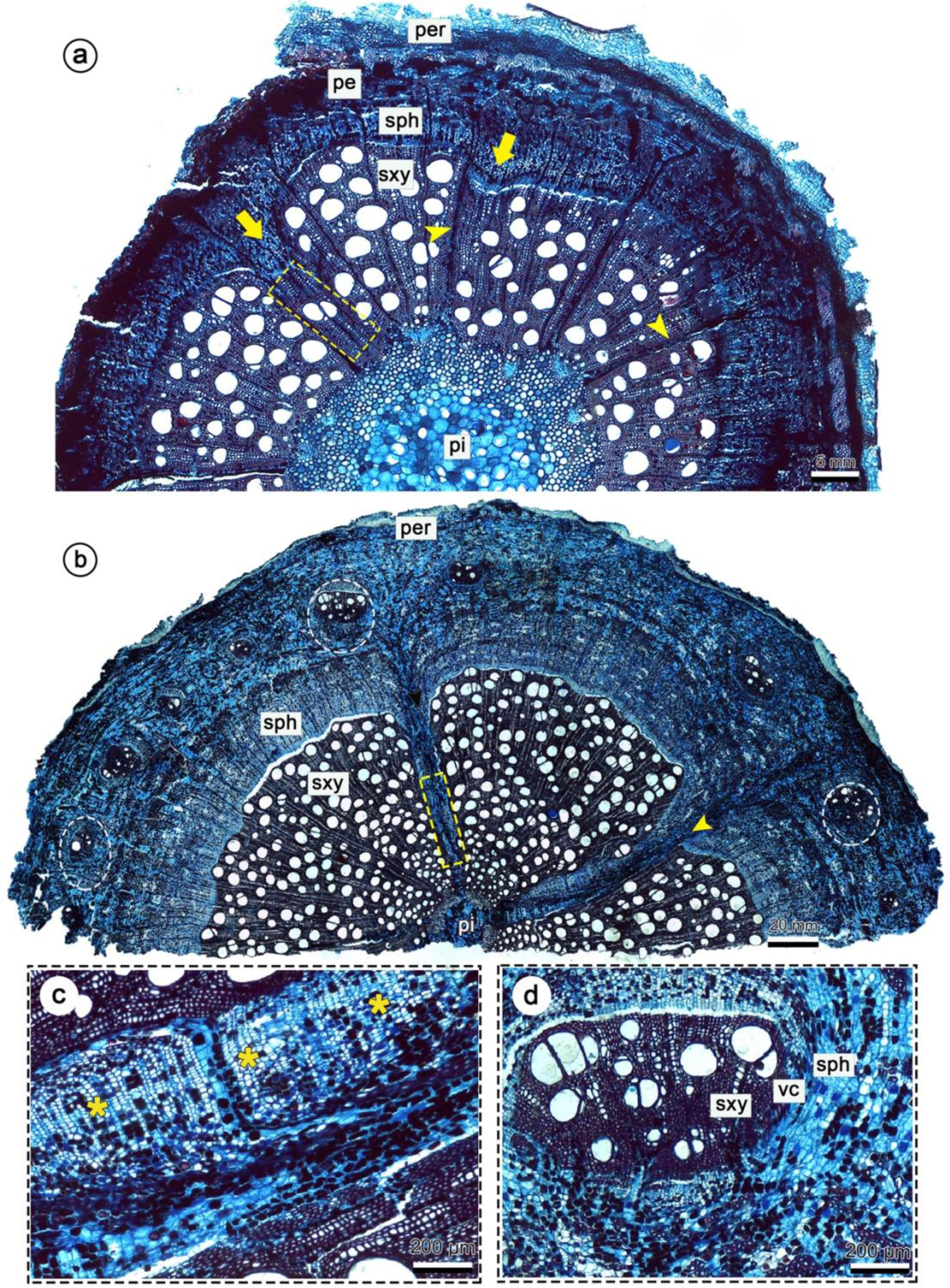
Development of fissured stems and ectopic cambia in *Serjania piscatoria* (individual 1). A, Stem with ca. 6 mm showing differential cambial activity forming phloem wedges (yellow arrows), reduced number of vessels in the phloem wedge region (yellow dashed rectangle), and wid vascular rays (yellow arrowheads). B–D, Stem with ca. 20 mm showing deep phloem wedges (pw) extending from the inner portion of the stem near the pith (yellow arrowheads) to the most external region of the secondary phloem; note the continuous cambium in the inner portion of the phloem wedge (C), and neoformations (ectopic cambia) arising in the non-conductive secondary phloem (D) (white dashed ellipses in B). Abbreviations: per = periderm, pe = pericycle, pi = pith, sxy = secondary xylem, sph = secondary phloem, vc = vascular cambium.

### Phloem arcs and isolated bundles in early developmental stages

During the transition from primary to secondary growth (approximately 5 mm in diameter), the stem has a nearly circular conformation, although there is a greater formation of secondary phloem compared to the secondary xylem giving rise to phloem arcs in several portions of the stem (Fig. 2A-C). These areas usually coincide with the region of the interfascicular cambium (Fig. 2A, C). With the continued differential activity of the cambium during stem development, these phloem arcs progress into phloem wedges, which are regions with a greater disparity in the ratio of phloem to secondary xylem (Fig. 2C). This process is an early step towards the formation of deep phloem wedges as will be discussed in the section “Differential cambial activity: the expansion of phloem wedges”.

In some individuals, one or two vascular bundles are isolated from the rest of the eustele bundles in young stems (approximately 1-2 mm in diameter) (Fig. 3A-D). These isolated bundles consist of primary xylem and phloem with the usual polarity, i.e., centripetal xylem and centrifugal phloem (Fig. 3B-E, 4A-C). In some cases, concurrent with cambial activity in the central cylinder (Fig. 3B-E, Fig. 4A-B), the bundle may remain isolated along a few internodes, and a cambium may develop from the procambium of the bundle (Fig. 3B-D, Fig. 4B). In other cases, this isolated vascular bundle from one internode above (Fig. 3D) merges with the central vascular cylinder after passing through the node and no longer appear completely isolated (Fig. 3E).

During secondary growth, specimens with an isolated bundle (approximately 7 mm in diameter) form a typical vascular cylinder derived from the eustele bundles in the central region, while an independent cambium forms secondary vascular tissues with the usual polarity around the “isolated bundle” (Fig. 4D-E). The cambium in the “isolated bundle” develops concentrically from the procambium of the bundle and determines the formation of a circular vascular unit, hereafter referred to as the “vascular unit” (Fig. 4D). The protoxylem pole can be observed at the center of the vascular unit (Fig. 4E), confirming its origin from the isolated vascular bundle. This vascular unit persists in the adult stems of these specimens (Fig. 4D).

### The expansion of phloem arcs into phloem wedges in mature stems

As secondary growth progresses (stems with ca. 6 mm), the vascular cambium of the central cylinder maintains a differential activity in the phloem arcs that emerged in early developmental stages, generating increasingly deeper wedges (Fig. 5A-C). The phloem wedges exhibit distinct sizes along the stem circumference (Fig. 5A). There is a lower production of vessel elements in the phloem wedges, with secondary xylem formed mostly by fibers (Fig. 5A). At this stage of development, the stem maintains a circular outline, with a continuous cambium (Fig. 5B), which persists even in the inner regions of the phloem wedge (Fig. 5C).

### Dissection of the vascular system and the emergence of ectopic cambia

In developed stems (approximately 20 mm in diameter), the differential cambial activity continues between the phloem wedges and the inter-wedges (Fig. 5D-E). The cambium remains continuous and there is a higher production of vascular tissues in the region outside the wedge compared to the vascular cambium in the phloem wedge region (Fig. 5D-E). This process leads to the formation of deeper phloem wedges that dissect the secondary xylem into various portions (Fig. 5D). Generally, three to four major phloem wedges are observed (Fig. 5D), while smaller wedges may develop in other regions (Fig. 5D). In parallel with the dissection of the vascular system by phloem wedges, ectopic cambia arise from the differentiation of axial parenchyma of the non-conducting secondary phloem (Fig. 5D-E). These new cambia can differentiate parallel or perpendicular to the stem axis (Fig. 5F). The new cambia continue to produce secondary xylem centripetally and secondary phloem centrifugally, tending to develop circular vascular units (Fig. 5F).

### The combination of ontogenetic processes in adult stems

Individuals in advanced secondary growth accumulate a series of anatomical modifications resulting from the combination of two or more of the four ontogenetic processes described earlier (Fig. 6A-H). In general, most adult individuals display a compartmentalized vascular system due to the formation of deep phloem wedges (Fig. 6 B-H) and the emergence of ectopic cambia (Fig. 6B-D; F-H). Due to the random reiteration of these processes, the mature stem assumes highly distinct and intricate vascular configurations in each specimen, characterizing the species’ polymorphism (Fig. 6G-H; Supporting Information, Fig. S2; Supporting Information, Video S1).

**Figure 6.**
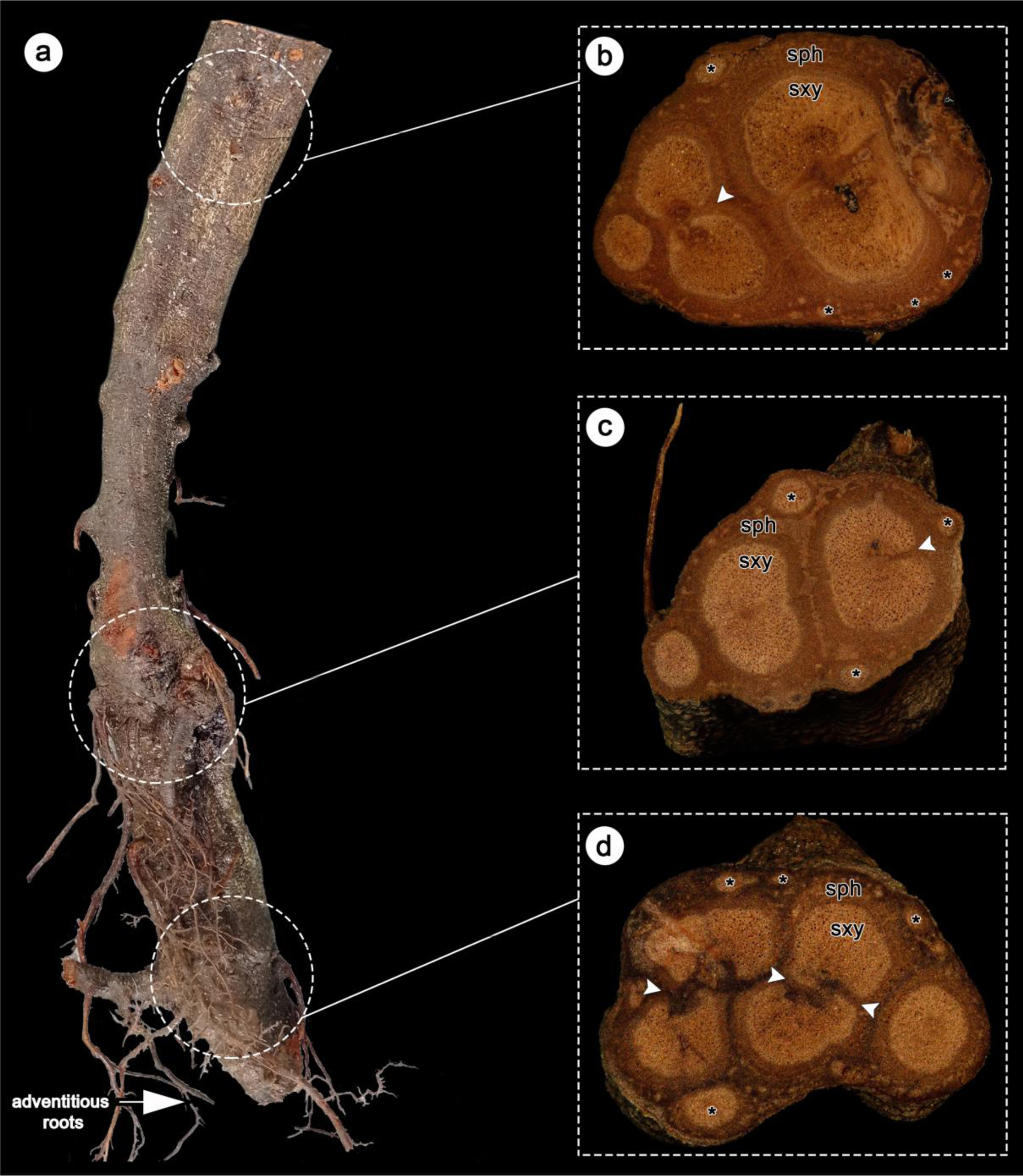
Macromorphology of adult stems of *Serjania piscatoria* (individual 3). A, Stem near the base of the plant. B–D Details of A showing cross-sections sampled at 150–200 mm away from the base; note the complex vascular system, resulting from multiple ontogenetic processes, including fragmentation by phloem wedges (white arrowheads) and neoformations (ectopic cambia) (asterisks). Images were photographed at intervals of approximately 50 mm. Abbreviations: sxy= secondary xylem, sph= secondary phloem. Stem diameters: Stem diameter: A = 20 mm, B = 20 mm, C = 37 mm, D = 52 mm.

### The phylogenetic position of *Serjania piscatoria*

The maximum likelihood (ML) phylogenetic tree (Fig.7A) and the 50% majority rule consensus Bayesian tree (Supporting Information, Fig. S3) are congruent but differ in the position of three species, which are *S. ichthyoctona* Radlk. (clado 8), *S. tortuosa* (Benth.) Ferrucci & V.W.Steinm. (clade 7) and *S. altissima* (Poepp.) Radlk. (clade 2) (Fig. 7A). The internal nodes have bootstrap support values ranging from 60 to 90 in the ML tree and posterior probability ranging from 0.7 to 1 in the Bayesian tree. The tree was rooted with *Thinouia*, with *Lophostigma* as the first lineage to diversify, followed by the genus *Urvillea* (Fig. 7A). *Cardiospermum* is polyphyletic, with some species forming a sister clade to *Paullinia*, and *C. urvilleoides* nested within *Serjania* (Fig. 7A). *Serjania piscatoria* is weakly supported as the sister lineage to *Serjania goniocarpa* Radlk., and they are nested within clade 5, along with seven other species (Fig. 7A): *S. membranacea* Splitg., *S. clematidifolia* Cambess., *S. fuscifolia* Radlk., *S. pyramidata* Radlk., *S. paucidentata* Radlk., *S. incana* Radlk., and *S. grandiceps* Radlk.

**Figure 7.**
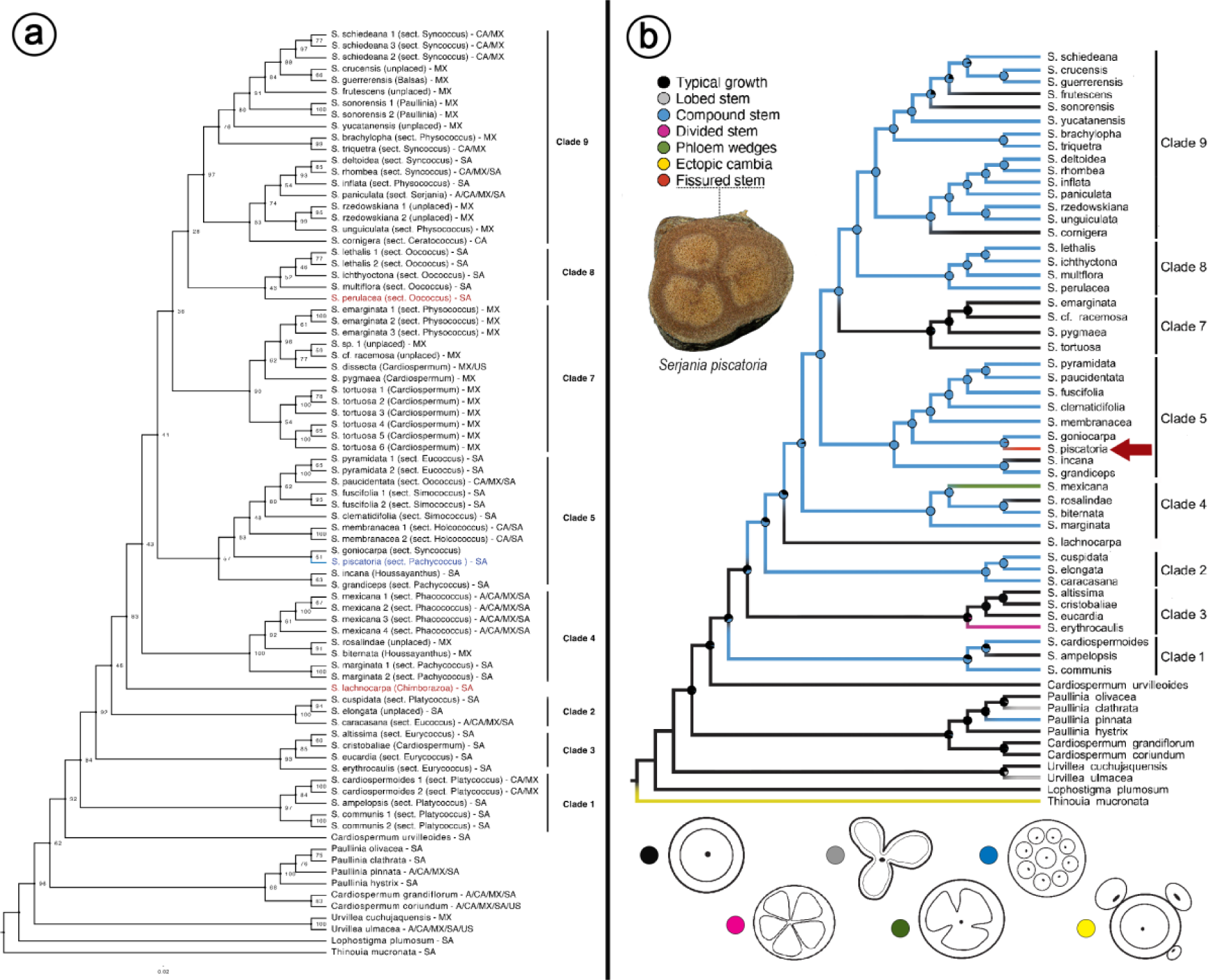
*Serjania* phylogeny indicating the position of *Serjania piscatoria* and ancestral state reconstruction of vascular variants in the genus. A, Maximum likelihood phylogenetic tree of *Serjania* showing *S. piscatoria* within clade 6 in blue; the numbers associated with the nodes indicate bootstrap values; clades are numbered following Steinmann et al. (2022), excluding clade 6 which was split in this analysis and are indicated in red. Taxon names are followed by geographic regions: Central America (CA), Mexico (MX), South America (SA), United States (USA), Antilles (A). B, Ancestral state reconstruction showing the pattern of evolution of vascular variants in *Serjania* and outgroups. The ancestor of *Serjania* was reconstructed with a higher probability of having a simple stem. Within *Serjania*, all vascular variants evolved only once in this analysis, including the fissured stem in *S. piscatoria.* Drawings for stem types adapted from Pace et al., 2022, modified with permission from Springer Nature (license 5750810420077).

### The evolution of vascular variants in *Serjania*

The ancestral state reconstruction indicated that the most recent common ancestor of Paullinieae and *Serjania* likely had typical growth, and compound stem evolved at least twice, in clade 1, and the lineage derived from the common ancestor of clades 2, 4, 5, 7, and 9 (Fig. 7B). In this analysis, *Serjania piscatoria* was coded as having a “fissured stem” (i.e., highly dissected vascular cylinder resulting from the modified activity of a single cambium), which is the most common pattern observed in most specimens of adult stems for this species (except for samples at the base of the plant). This pattern evolved only in this lineage from an ancestor with a compound stem, which is the pattern found in all other species in clade 5 and in most other *Serjania* species included in the phylogeny (Fig. 7B). The other patterns arose independently only once, namely: phloem wedges observed in *S. mexicana* (clade 4); divided stem – which generates vascular cylinders from the rise of independent vascular cambial segments in the lobes, with or without a central cylinder in old stems – observed in *S. erythrocaulis* Acev.-Rodr. & Somner (clade 3); lobed stem – characterized by lobes and furrows generating non-cylindrical stems – observed in *Paullinia clathrate* Radlk., and ectopic cambia observed in *Thinouia mucronata* Radlk. (Fig. 7B). Additionally, there were multiple reversals from compound type to typical stem within *Serjania*, as in clades 4, 5, 7, and 9.

## DISCUSSION

### *Serjania piscatoria*: a case of developmental lability and polymorphism

Stem development in *S. piscatoria* involves four major developmental processes leading to atypical vascular architectures that diverge from the typical pattern observed in seed plants. These four ontogenetic processes can be classified into the three categories of vascular variants (Cunha Neto, 2023). This anatomical variability characterizes *S. piscatoria* as a case of labile stem development, as well as a polymorphic species given that these different processes are not always present in all individuals of the same population. Below we discuss each of these ontogenetic processes in detail and their implications for the diversity and evolution of the vascular system in Sapindaceae lianas.

### The isolated bundle: the path to “alternative” compound stems

A remarkable phenomenon in the stem development of *S. piscatoria* is the alternative presence of an isolated vascular bundle during primary growth, which can persist independently in the adult stem, forming a vascular unit with secondary growth. Given its origin from the procambium during primary growth, this phenomenon is classified as a “procambial variant”, the first category of vascular variants by Cunha Neto 2023. This phenomenon, which was mentioned by Schenck (1893), is also reported for the species *Serjania subdentata* Juss. ex Poir., characterized as having a compound stem (Johnson and Truscott, 1956). In that study, the authors demonstrated that one or more bundles can remain separate from the eustele and may fuse permanently or temporarily to the central cylinder after passing through some nodes of the stem (Johnson and Truscott, 1956). In both *S. piscatoria* and *S. subdentata*, if one of these bundles remains isolated, a cambium is formed and produces secondary vascular tissues, giving rise to a circular (isolated) vascular unit. An isolated vascular bundle giving rise to a vascular unit with secondary growth is also the process described for *Paullinia pinnata* L. (Van der Walt, 1973; Chery et al., 2020), a species traditionally described with a compound stem. In species compound stems, the vascular units in the secondary body (derived from independent/isolated bundles) form the so-called peripheral vascular cylinders (Cunha Neto, 2023). Since the development among these different species proceeds from the same ontogenetic process, the stem of *S. piscatoria* can also be classified as having a compound stem, in this case, considered facultative, as this structure is not always present across specimens. Compound stems derived from a single isolated bundle have always been associated with the genus *Paullinia* (based on *P. pinnata*; Van der Walt, 1973; Chery et al., 2020; Cunha Neto, 2023). However, studies on *S. subdentata* and *S. piscatoria* reveal that this phenomenon is also present in *Serjania*.

### Phloem wedges: the path to fissured stems

The formation of phloem arcs progressing into phloem wedges and deep invaginations that ultimately compartmentalize the vascular system (cambial variant) is another phenomenon observed in stems of *S. piscatoria*. Given its origin from the cambium during secondary growth, this phenomenon is classified as a “cambial variant”, the second category of vascular variants by Cunha Neto 2023. It is no longer surprising that Schenck (1893) had already made observations in this regard. What was unclear is that, in a perspective of the evolution of development, phloem wedges can be both the final stage of development and an intermediate step towards more complex anatomical patterns, which is the case also of *S. piscatoria*. In the first case, adult stems exhibit only “phloem wedges”, a pattern that shares the same name as the ontogenetic process and is reported to some Paullinieae (e.g., *U. stipularis* Ferrucci; Cunha Neto et al., 2023), and species in other families (e.g., Bignoniaceae; Pace et al., 2009). In the second case, fissured stems emerged from lineages containing phloem wedges, supporting the latter pattern as an intermediate step in the formation of more complex morphotypes. This scenario has been reported for both Paullinieae (e.g., *Urvillea*; Cunha Neto et al., 2023) and other families (e.g., Malpighiaceae; Quintanar-Castillo and Pace 2022). Unlike Malpighiaceae (Cabanillas et al., 2017; Quintanar-Castillo and Pace, 2022), the proliferation of parenchyma contributing to the dissection of secondary xylem in fissured stems was not observed in *S. piscatoria*.

### Ectopic cambia: another path to structural diversity

Similar to other Paullinieae, *S. piscatoria* presents ectopic cambia (Tamaio and Angyalossy, 2009; Tamaio and Somner, 2010; Cunha Neto et al., 2018, 2023; Rajput et al., 2021, Rizzieri et al., 2021), which is the third category of vascular variants by (Cunha Neto, 2023). In the stem of *S. piscatoria*, these new cambia frequently generate circular units irregularly distributed along the circumference of the stem (in addition to eventual peripheral cylinders derived from the isolated vascular bundle). This pattern is a typical case of neoformations – new cambia forming independent, circular vascular units in an irregular fashion (Cunha Neto and Onyenedum, 2023) – similar to that described for stems (Tamaio and Angyalossy, 2009; Cunha Neto et al., 2018; Rizzieri et al., 2021) and roots (Bastos et al., 2016) of other Paullinieae species, and other families (e.g., Rubiaceae, Leal et al., 2020). In both *S. piscatoria* and other Paullinieae species, neoformations are produced after the establishment of other patterns of vascular variants, including successive cambia (Cunha Neto et al., 2018), which is also a type of ectopic cambia, as well as divided stems (Rizzieri et al., 2021), and compound stems (Tamaio and Angyalossy, 2009), which are cases of procambial variants (Cunha Neto 2023). Regarding their origin, neoformations arise from vascular parenchyma in Paullinieae, but in lianas from other families, they may originate from the cortex (e.g., Rubiaceae, Leal et al., 2020). Interestingly, neoformations (=ectopic cambia) was likely present in the first fossil of Sapindaceae, i.e., *Ampelorhiza heteroxylon* gen. et sp. nov. (Jud et al., 2021). In this specimen, some of the hypothetical “peripheral cylinders” may represent neoformations (originating from a new cambium) given their small size compared to the remaining vascular cylinders. These observations indicate that the ability to form ectopic cambia was already present in Paullinieae species since the Miocene.

### *Serjania piscatoria* in the genus phylogeny

Overall, our phylogenetic hypothesis is similar to the topology obtained by Steinmann et al. (2022), and both of them are consistent with other phylogenies within Paullinieae (Chery et al., 2020, Medeiros et al., 2020; Cunha Neto et al., 2023). These three studies focus on the genera *Paullinia*, *Thinouia*, and *Urvillea*, respectively, using representatives of the other five genera of Paullinieae as outgroups. In most studies, *Thinouia* emerges as the sister lineage to the other genera of Paullinieae (Acevedo-Rodriguez et al., 2017; Medeiros et al., 2020; Chery et al., 2020; Cunha Neto et al., 2023) and this genus was used to root the tree in the present study. Following this approach, the position of *Lophostigma* and *Urvillea* are altered compared to the original tree presented by Steinmann et al. (2022). Furthermore, with the inclusion of *S. piscatoria,* clade 6 of Steinmann et al. (2022) was dissolved, with one of the two species of this clade grouping into clades 8 (*S. perulacea* Radlk.) and the other appears as the sister lineage of clades 4, 5, 7, and 8 (*S. lachnocarpa* (Benth. ex Radlk.) Acev.-Rodr.).

*Serjania piscatoria* grouped with *Serjania goniocarpa* within clade 5. The support for this relationship was low (= 51 bootstrap), which is not significantly different from values found for some clades within the phylogeny by Steinmann et al. (2022). The low support in these analyses can be probably explained by the small sampling relative to the size of the genus, which contains around 270 species (World Flora Online, 2023). It is important to note that *S. piscatoria* and *S. goniocarpa* do not exhibit unique morphological characteristics that justify their grouping, as indicated by stem and leaf characteristics (Supporting Information, Table S1). The same applies to the other species in clade 5. Additionally, *Serjania piscatoria* and *S. goniocarpa* have disjunct distributions (although within the same biogeographical region: South America including Central America and the West Indies; Buerki et al., 2011), with the former being restricted to the Atlantic Forest in southeastern Brazil, and the latter occurring in Central America (Belize, Guatemala, Honduras, and Mexico; https://www.gbif.org/). These observations point to the need for further examination of morphological and molecular studies to enhance the resolution of the *Serjania* phylogeny, as well as the reconstruction of biogeographical scenarios within Paullinieae.

### Initial evidence on the evolution of vascular variants in *Serjania*

As mentioned earlier, *S. piscatoria* stands out for having a highly labile stem development. Studies on the evolution of vascular variants typically characterize species by the presence or absence of the studied developmental pathways, culminating with a single anatomical pattern (Chery et al., 2020; Cunha Neto et al., 2023). To conduct a similar analysis in the ancestral state reconstruction, we coded *S. piscatoria* as having a “fissured stem”, as this pattern is derived from ontogenetic modifications that begin in early secondary growth and because it is present in most of the studied specimens/individuals. The ancestral state reconstruction indicated that the fissured stem evolved from an ancestor with a compound stem. Biologically, this scenario is unexpected, given that the compound stem is formed by modifications in the primary vascular system (Tamaio and Angyalossy 2009). In this context, the evolution of fissured stems from compound stems imposes numerous changes in development, from the reversion to the typical eustele to the emergence of a single cambium with differential activity forming phloem wedges, which are a pre-requisite to this type of compartmentalization of the vascular system. On the other hand, the emergence of the “facultative compound” pattern in *S. piscatoria* is more easily explained in this context, since all other species in this clade exhibit compound stems, which may have facilitated the maintenance of this phenomenon in *S. piscatoria*. Schenck (1893) hypothesized that the compound structure would have evolved from the typical stem, where the “first type” of compound stems to evolve would have three peripheral vascular cylinders. In this context, the author points to *S. piscatoria* as an interesting species in the evolution of compound stems, as it presents an indefinite number of “peripheral cylinders,” ranging from one to three cylinders. Based on our results, what Schenck called “peripheral cylinders’’ comprise structures derived from different processes including the “isolated vascular unit” (i.e., facultative compound) and neoformations, which is derived from *de novo* cambia. The facultative compound structure with one or two peripheral vascular cylinders observed in *S. piscatoria* is developmentally similar to the compound stem reported to *S. dentata* (Johnson and Truscott, 1956), which is also described to *Paullinia* species (var der Walt 1973; Chery et al., 2020). Nevertheless, in most *Serjania,* compound stems typically derive from multiple pre-existing bundles (including some with inverted polarity) and may form up to nine or ten peripheral vascular cylinders (Tamaio and Angyalossy, 2009). This indicates that compound stems may be formed by at least two different developmental programs in Paullinieae lianas (Pace et al., 2022; Cunha Neto 2023) and that they are formed constitutively in these species, which may have involved an intricate diversification in the number and organization of stem vascular bundles through evolutionary time. Future studies of the evolution of development expanding species sampling including other known patterns for the genus and the tribe (e.g., typical; divided; lobed) would shed light on the pattern of evolution of vascular variants in the group.

From a developmental and evolutionary perspective, the evolution of ectopic cambia in *S. piscatoria* is not surprising. Ontogenetically, it is known that this phenomenon occurs in species of different genera of Paullinieae, either as a single type of vascular variant (e.g., successive cambia, *S. pernambucensis* Radlk., and some *Paullinia* species; Cunha Neto et al., 2018) or in combination with other variants, as observed with the emergence of neoformations in species with divided (Rizzieri et al., 2021) and compound stem (Tamaio and Angyalossy, 2009). Previous studies on the evolution of development have shown that ectopic cambia evolved only once in *Paullinia* (= successive cambia; Chery et al., 2020) and once in *Urvillea* (Cunha Neto et al., 2023). In the case of *Serjania*, ectopic cambia – the developmental capacity to naturally generate new cambia – likely evolved multiple times, as different patterns are observed, including corded stems (e.g., *S. meridionalis* Cambess; Borniego and Cabanillas, 2014), successive cambia (e.g., *S. pernambucensis*; Cunha Neto et al., 2018), and neoformations (e.g., *S. caracasana* (Jacq.) Willd.; Tamaio and Angyalossy, 2009). However, several species are not yet represented in the present phylogeny, pointing to the need for larger investigations of the evolution of development within Paullinieae to allow more accurate analysis of the pattern of evolution of these complex phenotypes.

## CONCLUSIONS

We demonstrate the interaction of developmental processes generating the stem vascular architectures of *S. piscatoria*, which include modifications arising at the procambium, cambium, and ectopic cambia levels. The existence of multiple developmental processes represents a significant lability of vascular meristems and a polymorphism characteristic of *S. piscatoria* (although other examples of polymorphic species might exist within the genus, e.g., *S. multiflora* Cambess; “Lianas and Climbing Plants of the Neotropics” (Acevedo-Rodríguez et al. 2015 onwards). The polymorphism of *S. piscatoria* is a reminder of the great stem morphological diversity of Paullinieae lianas, and that the use of this trait for systematics should carefully observe this variability. Such cases add up to the prominent confusion between vascular variants forming a cable-like structure (e.g., compound, divided, corded stems), where ontogenetic modifications may occur either in early or later developmental stages creating stems with similar morphotypes though originated from distinct developmental processes. This study highlights the importance of developmental investigations to disentangle the mechanisms underlying the diverse vascular architectures observed in Paullinieae lianas and contributes to its systematics through the inclusion of *S. piscatoria* in the *Serjania* phylogeny for the first time. In addition, ancestral state reconstruction indicates that the vascular architecture observed in *S. piscatoria* evolved within a clade of species with compound stems, raising new questions on the evolution of vascular variants for the group. Future studies will be necessary to unravel more clearly the details of the time and mode of evolution of this phenomenon within the large and charismatic Paullinieae tribe, which holds the largest diversity of vascular variants in the plant kingdom.

## SUPPLEMENTARY DATA

Supplementary data will be available online.

**Figure S1.** Stem of individual 4 sampled from 100 mm away from the base of the plant, showing images of cross-sections at intervals of ca. 50 mm.

**Figure S2.** Stem of individual 3 sampled from 300-500 mm away from the base of the plant, showing images of cross-sections at intervals of ca. 50 mm.

**Figure S3.** Bayesian 50% majority-rule consensus tree from a Bayesian analysis of the combined, two-marker dataset for *Serjania* and outgroups.

**Dataset S1.** Dataset for ancestral state reconstruction of vascular variants.

**Table S1.** Comparison of morphological characters between *Serjania piscatoria* and *S. goniocarpa*.

## ACKNOWLEDGMENTS

This work is part of the master’s dissertation of the first author, who acknowledges the use of Google Translate (Google, https://translate.google.com/) for translation from Portuguese to English, and ChatGPT 3.5 (Open AI, https://chat.openai.com) to support proofreading of the initial draft. Grammarly (Grammarly Inc., https://grammarly.com) was used for final proofreading of the manuscript. We thank Dra. Neusa Tamaio and Carolina L. Bastos for their help providing plant samples and images. We also express our gratitude to the reviewers who contribute to scientific advancement without any compensation.

## AUTHOR CONTRIBUTIONS

All authors contributed to the design, analysis, writing, revision, and proofreading of the final version of the manuscript.

## DATA AVAILABILITY

The DNA sequence alignment, data for ancestral state reconstruction, and a video illustrating a specimen with vascular variants are available via Zenodo (https://zenodo.org/records/10821731).

