## Supplementary figures and images for "The “abominable mystery” of Schenck: the polymorphism of *Serjania piscatoria* and its implications for the evolution of vascular variants in Paullinieae (Sapindaceae)"

### Figure_S1.tif

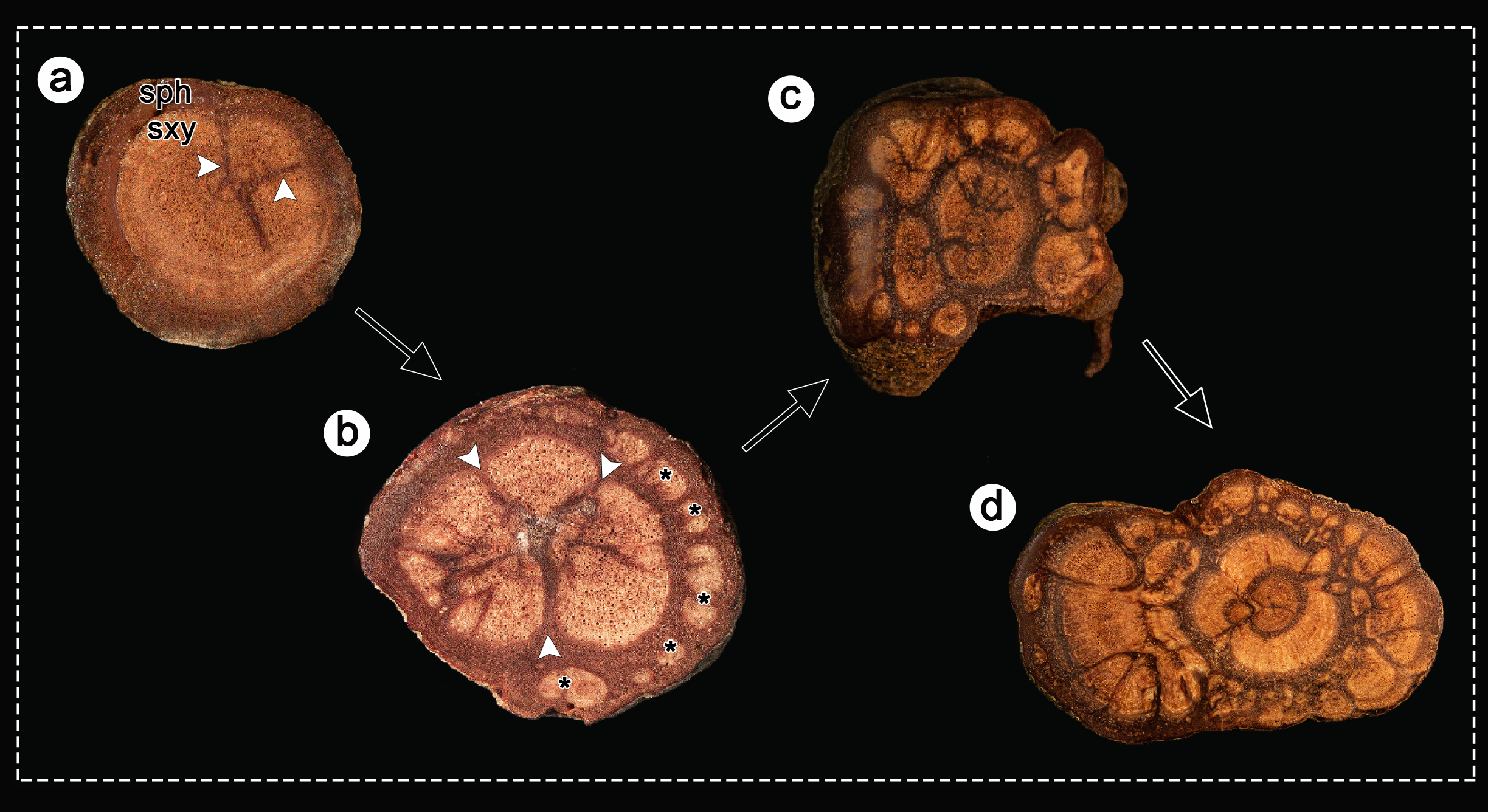

### Figure_S2.tif

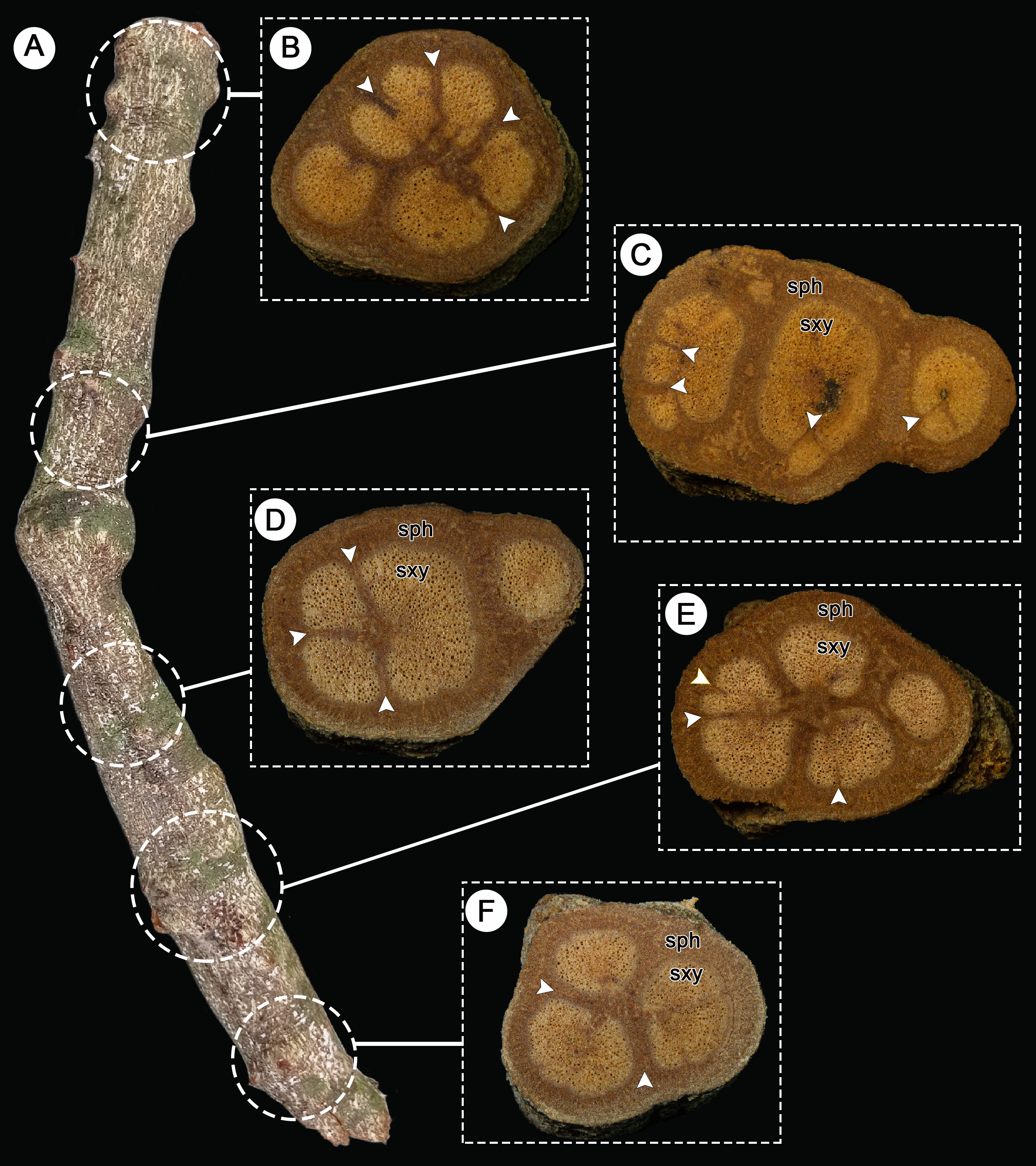

### Figure_S3.jpg

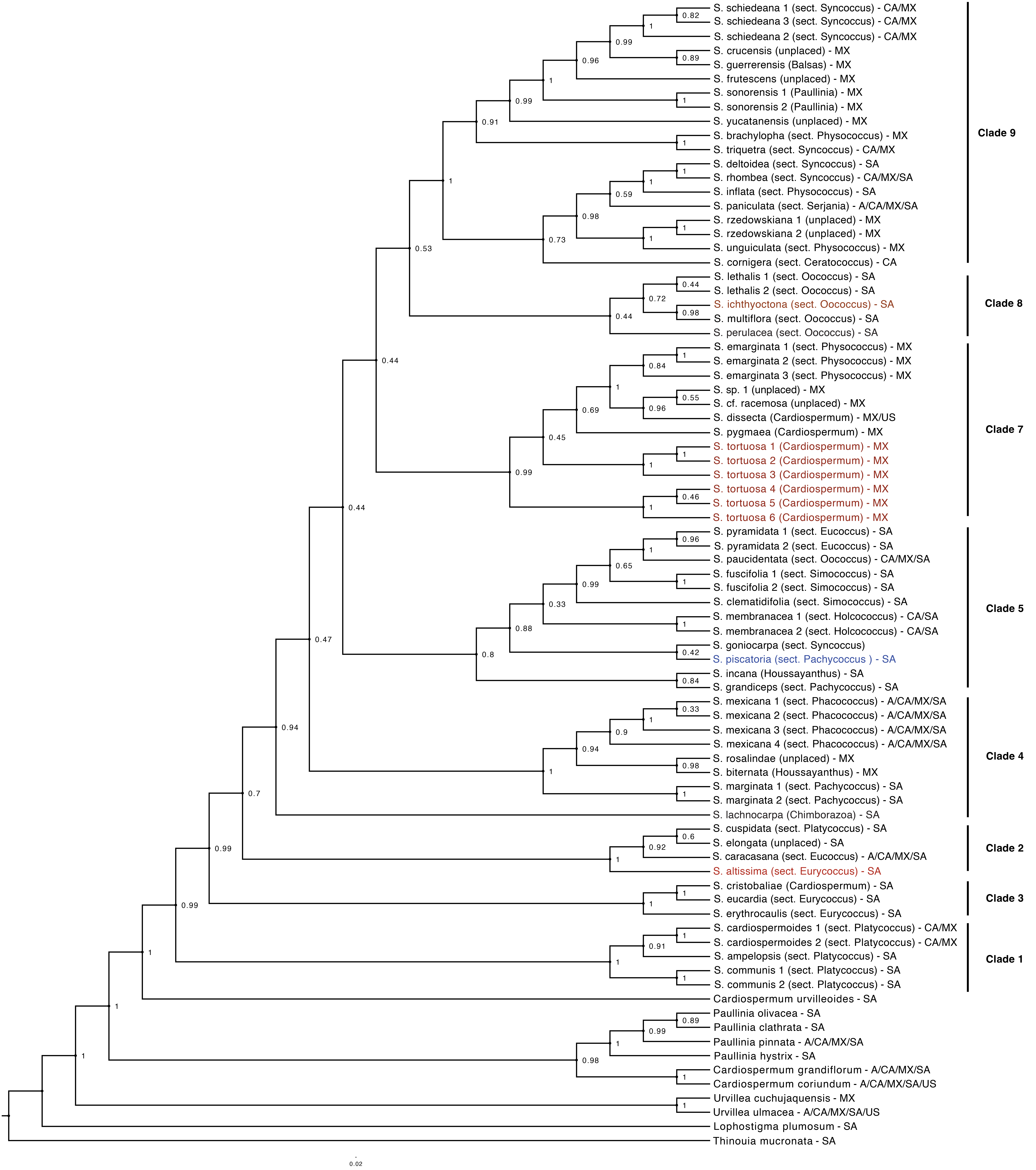
